# What can New Zealand bats tell us about Coronaviruses?

**DOI:** 10.1101/2023.05.26.542035

**Authors:** Pablo Tortosa, Kate McInnes, Colin F. J. O’Donnell, Moira Pryde, Yann Gomard, Camille Lebarbenchon, Robert Poulin

## Abstract

The current Covid-19 pandemic emphasizes the dramatic consequences of emerging zoonotic pathogens and stimulates the need for an assessment of the evolution and natural cycle of such microbes in a One Health framework. A number of recent studies have revealed an astonishing diversity of bat-borne Coronaviruses, including in insular environments, which can be considered as simplified biological systems suited for the exploration of the transmission cycles of these viruses in nature. In this work, we present two new lineages of alpha Coronaviruses detected by screening the only two extant New Zealand bat species: the lesser short-tailed bat (*Mystacina tuberculata*) and the long-tailed bat (*Chalinolobus tuberculatus*). Infection prevalence reaching 60% in long-tailed bats makes this host-pathogen model relevant for the investigation of maintenance mechanisms in a bat reservoir with peculiar physiological adaptations to temperate climates. A phylogenetic analysis shows that these viral lineages do cluster with Coronaviruses hosted by bat sister species from Australia, supporting co-diversification processes and confirming that the evolution of these viruses is tightly linked to that of their hosts. These patterns provide an interesting framework for further research aiming at elucidating the natural history and biological cycles of these economically-devastating zoonotic viruses.

## Introduction

The current Covid-19 pandemic has brought to the fore front investigations on zoonoses and further illustrated the importance of understanding both the evolutionary history and the natural cycles of infectious agents with zoonotic potential. While an animal origin for human Coronaviruses (CoVs) is reaching consensus (Chan et al., 2013; Cui et al., 2019), the actual identification of the species acting as reservoirs for such emergent viruses is still far from complete. Further, the natural transmission cycles of these viruses remain essentially a black box, stressing the need for a comprehensive characterization of biotic and abiotic parameters influencing viral transmission in nature with the aim of highlighting putative emergence drivers.

Considering the high diversity of bat species worldwide, current knowledge on bat-borne Coronaviruses remains limited and strongly biased by oversampled taxa or regions (Anthony et al., 2017). For example, some tropical regions such as South East Asia (Lacroix et al., 2017), Western Africa (Maganga et al., 2020) or the islands of the Southwestern Indian Ocean (Joffrin et al., 2020) have been extensively explored while several resource limited countries have been hardly investigated (Anthony et al., 2017). Given that Chiroptera is the second most diversified mammalian order following Rodentia, the actual diversity of bat-borne Coronaviruses is certainly underestimated, especially considering that each bat species may host an average of 2.5 distinct viral lineages (Peel et al., 2020). The diversity of bat-borne Coronaviruses and the apparent complexity of their evolutionary history, including co-diversification and host switching events (Anthony et al., 2017), make this host-pathogen system a relevant model for the investigation of the evolution of zoonotic viruses. Recent investigations have reported strong co-phylogenetic signals between bats and hosted Coronaviruses, which further strengthens the role of bats as natural reservoirs for these emerging pathogens (Joffrin et al., 2020; Latinne et al., 2020). Insular ecosystems do facilitate the investigation of these host-pathogen interactions as they exhibit some peculiar properties such as geographical isolation that limits migration of animal reservoirs and associated viruses, a limited number of species facilitating the screening of putative reservoirs, and the age of island formation that can be considered as time boundaries for divergence/co-divergence processes (Tortosa et al., 2012).

In the present report, we explore the diversity of bat-borne Coronaviruses in the relatively simple insular system of New Zealand. Indeed, the country hosts only two extant bat species, *Mystacina tuberculata* (Mystacinidae) and *Chalinolobus tuberculatus* (Vespertilionidae), known as the lesser short tailed- and long tailed-bat, respectively. Both are temperate rain-forest species, which primarily roost in colonies in tree cavities and rarely caves (O’Donnell & Sedgeley, 2006). Both species were common throughout the country, which, up until c. 800 years ago was fully forested. Over 75% of forests have been cleared since the arrival of humans in New Zealand, and both species are now threatened (O’Donnell et al., 2018; O’Donnell et al., 2010). However, both species are still widely distributed throughout the country, albeit in relatively low numbers and largely confined to forest remnants (Figure 1). Social organisation of the two species differs, with long-tailed bats forming adjacent small colonies, usually <200 bats (O’Donnell & Sedgeley 2006). In comparison, lesser short-tailed bats form colonies of >6000 individuals (Lloyd, 2001). Neither extant species are known to share colonial roosts and today, there are few locations where both species occur in the same forests.

**Figure 1:**
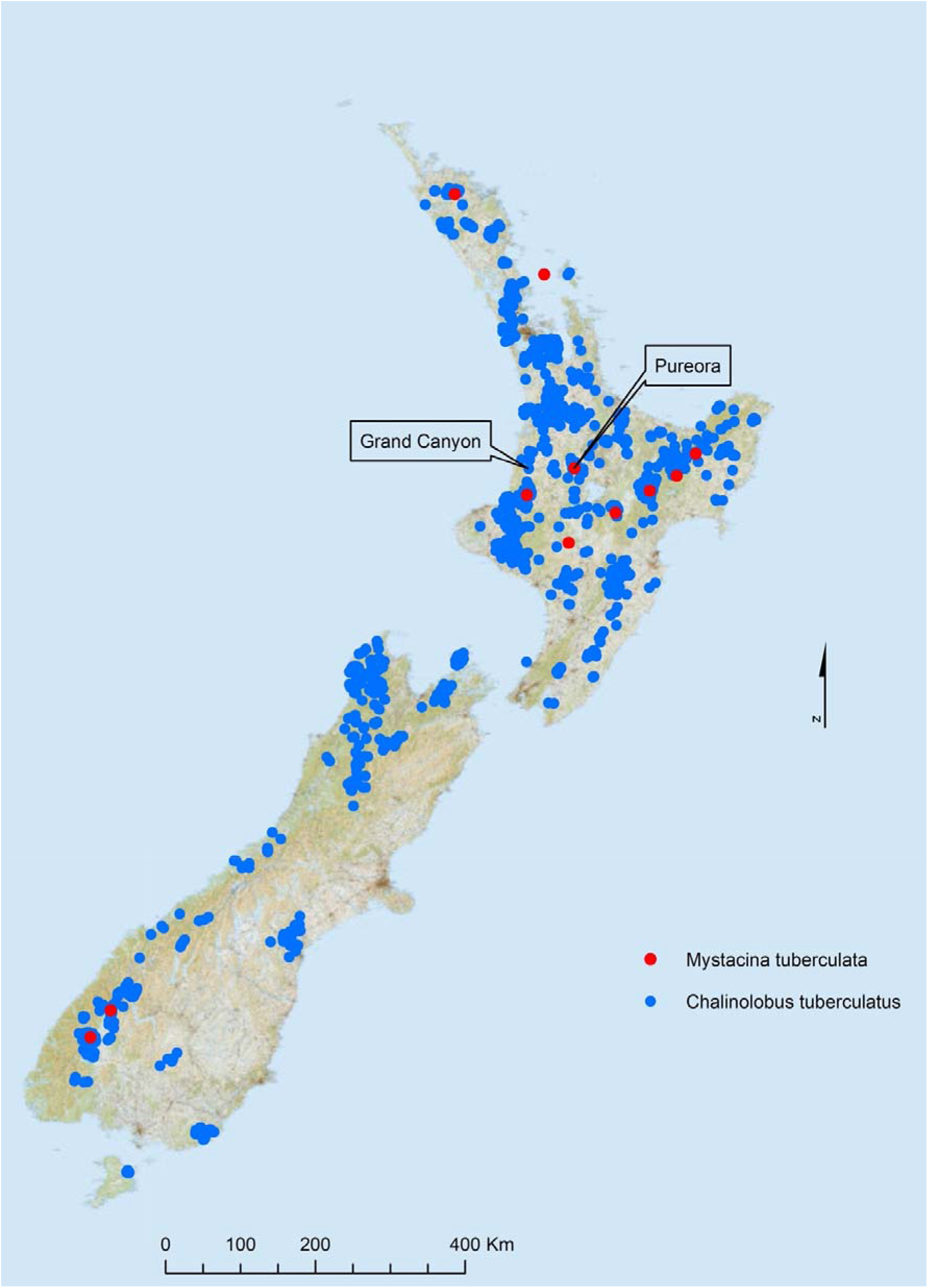
Occurrence of *C. tuberculatus* and *M. tuberculata*, and localization of the two sampling sites. Bat records include surveys done by DOC, councils, contractors, private groups, and individuals since 2002.

A previous study reported the full genome sequence of an alpha coronavirus obtained through massive sequencing of five bat scats from lesser short-tailed bats (Wang et al., 2015). We screened scats from 181 bat specimens belonging to both species and report molecular sequences revealing two new distinct CoV lineages hosted by New Zealand bats. The prevalence of CoV positive bats together with a phylogenetic analysis including CoVs hosted by Australian bat sister species pave the way for future studies aiming at inferring the evolution of these viruses as well as their natural cycle in a non-tropical insular environment.

## Methods

### Bats sampling

Lesser short-tailed bats (*M. tuberculata*) were sampled by collecting scats from clean tarpaulins that were spread at the bottom of tree holes used as day roosts near Pureroa, North Island (Figure 1). Scats were sampled the next morning using sterile toothpicks and stored individually in 500μL RNA later (Qiagen, Valencia, USA). Long-tailed bat (*C. tuberculatus*) specimens were trapped using harp traps in Great Canyon cave (Figure 1) and maintained individually in cotton bags from which one single scat was sampled and preserved in 500μL RNA later. Scats were then stored at room temperature and stored within a week following sampling in -70°C freezers until molecular analyses. All samplings were carried out by the New Zealand Department of Conservation staff, in conformity with the Conservation Act 1987.

### CoV screening

Nucleic acids were obtained using trizol (Invitrogen, Carlsbad, USA) extraction. Briefly, roughly half of each bat scat was taken out of RNA later using a sterile tip, and crushed into 1mL of cold trizol using a sterile pestle. Then RNA was extracted essentially following the manufacturer’s protocol and eventually resuspended in 40 μL RNAse-free water. cDNAs were obtained using Takara Primescript Reverse Transcriptase (Takara, Kusatsu, Japan) following manufacturer’s protocol. The detection of CoV cDNA was assessed using a previously published semi-nested PCR targeting a portion of the RNA-dependent RNA Polymerase (RdRp) encoding gene (Poon et al., 2005) except that the reverse primer was replaced by IN7* primer (5’-TAR CAM ACA ACA CCR TCA TCA GA-3’), which was designed using the numerous full CoV genomes made available since the original publication. Twenty samples randomly selected among the 50 Coronavirus positive samples (including the only two positive samples from short-tailed bats) reported herein were further Sanger sequenced on both strands at the Genetic Analyses Services from the University of Otago. CoV sequences are accessible in GenBank under accession numbers OK094715, OK094716 and OK094717.

### Phylogenetic analyses

We constructed a first Bayesian phylogeny essentially by using RdRp sequences included in previously published phylogenies presenting the diversity of Australasian bat-borne CoVs (Peel et al., 2020; Prada et al., 2019; Smith et al., 2016). JModelTest v.2.1.4 (Darriba et al., 2012) was used to determinate the best sequence evolution model. Then the Bayesian inference analyze was realized using Mr. Bayes v.3.2.3 (Ronquist et al., 2012) and GTR+I+G as substitution model with two independent runs of four incrementally heated Metropolis Coupled Markov Chain Monte Carlo starting from a random tree. The Metropolis Coupled Markov Chain Monte Carlo was run for 2 million generations with trees sampled every 100 generations. The convergence level of each phylogeny was verified by an average standard deviation of split frequencies inferior to 0.05. The initial 10% of trees for each run were discarded as burn-in and the consensus phylogeny with posterior probabilities were obtained from the remaining trees. The resulting phylogeny was visualized with FigTree v.1.4.2 (Rambaut, 2014).

We then extended the dataset by aggregating data presented in three published papers (Joffrin et al., 2020; Lacroix et al., 2017; Latinne et al., 2020), leading to a dataset of 1,150 sequences. Duplicated sequences as well as sequences that did not correspond to the RdRp locus were manually discarded from the dataset, leading to a final dataset composed of 812 sequences (see supplemental fasta file). An unrooted ML phylogeny was constructed using FastTree/One click on NGPhylogeny (Lemoine et al., 2019), which can be visualized on https://itol.embl.de/tree/92130128120274381648222266.

## Results and discussion

The study reveals high CoV infection prevalence (>60%) in long-tailed bats and limited infection in lesser short-tailed bats (<5%, see Table 1). Such figures suggest an important role of long-tailed bats in the maintenance of the detected CoV, but the limitations of the sampling scheme call for cautious interpretations. Strong temporal dynamics have been previously reported in CoVs transmission within bat colonies and the present data represent a single time point (Drexler et al., 2011; Joffrin et al., 2022; Wacharapluesadee et al., 2018). It is also important to note that long-tailed bats were individually processed by hand, identified at the species level and scats rapidly stored in RNA later. By contrast, scats from lesser short-tailed bats were obtained through a non-invasive method, and the absence of handling of each individual bat precludes definitive identification of these bat samples.

**Table 1:**
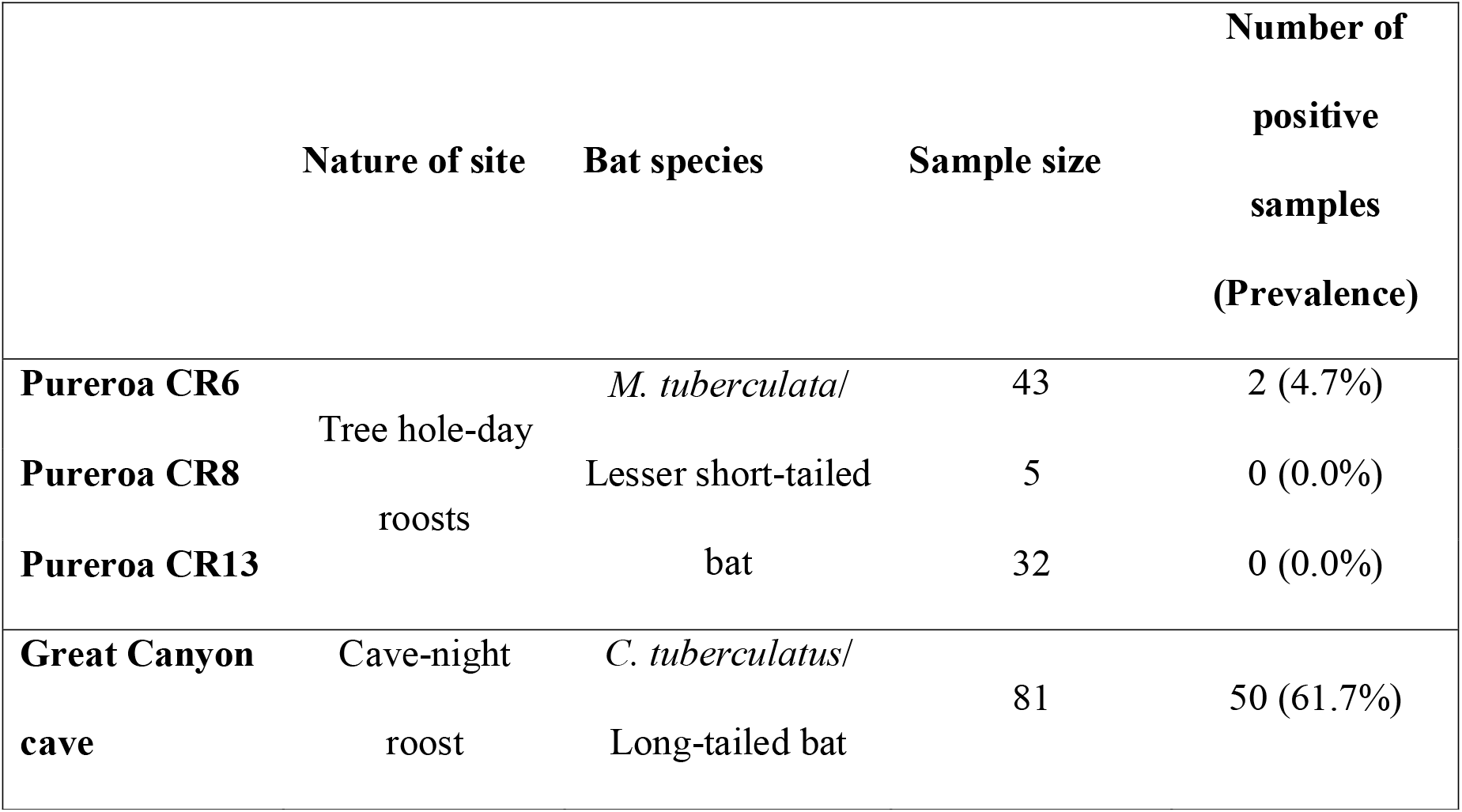
Sampled sites, species, and number of tested and coronavirus positive bats.

The phylogenetic analysis (Figure 2) reveals two distinct lineages with lineage 2 overwhelmingly dominant (19 out of the 20 obtained sequences, including the 2 sequences obtained from *M. tuberculata*). In addition, there is limited diversity within lineage 2, in keeping with previous studies showing limited intra-species diversity at the RNA-dependent RNA polymerase encoding gene (Joffrin et al., 2020). Although only Great Canyon cave was sampled for long tailed bats, opportunities for these bats to mix with individuals from unrelated colonies are likely high compared to lesser short-tailed bats (O’ Donnell et al., 2016; O’Donnell, 2000): long-tailed bats form temporary social groupings at night roosts, including at this sampling site where over 350 animals can appear at night, with the majority gone by morning (O’donnell, 2002).

**Figure 2.**
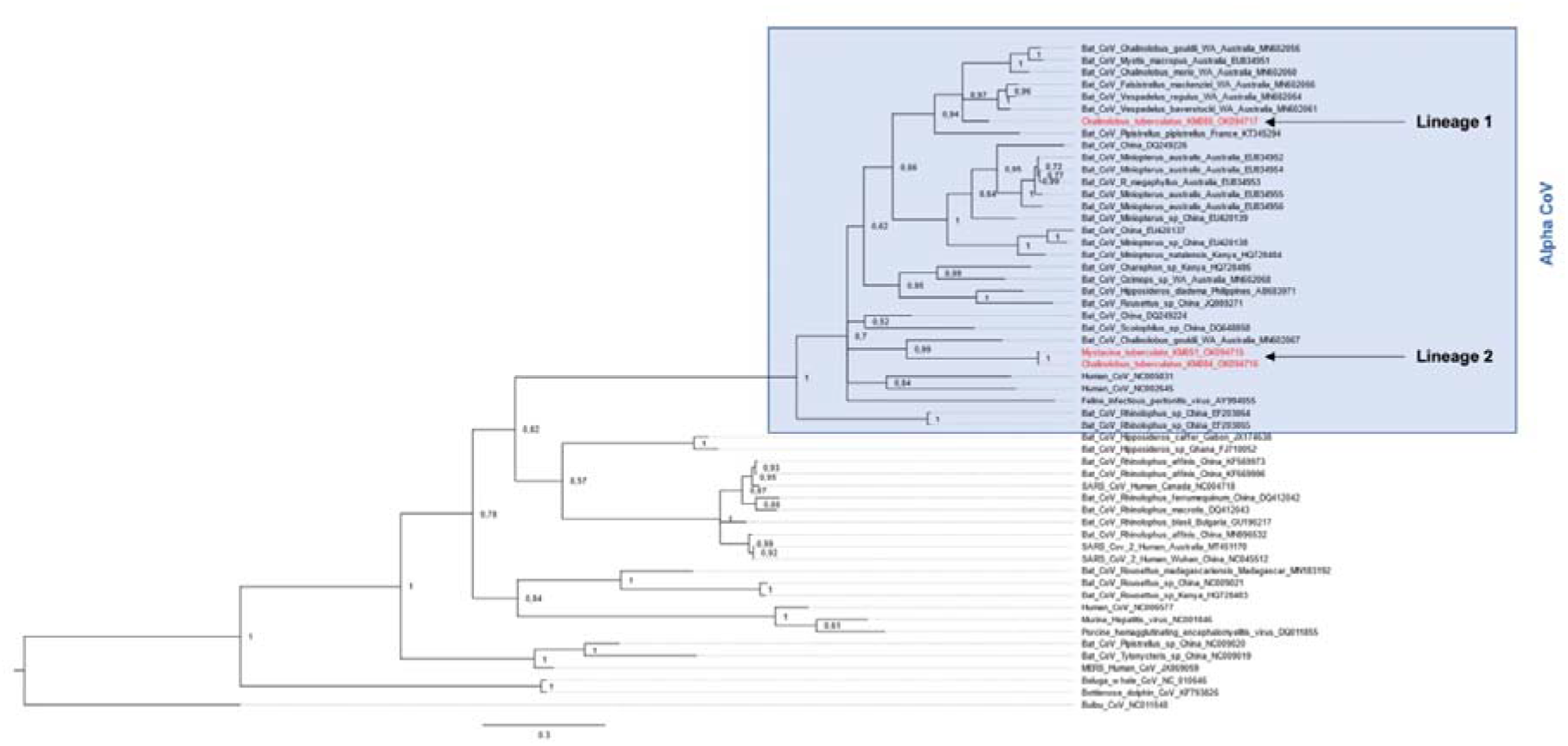
Bayesian phylogenetic reconstruction of bat-borne Coronaviruses using RdRp gene (406nt). Alphacoronaviruses are boxed in blue, lineages reported herein are highlighted in red. Adapted from (Peel et al., 2020).

Importantly, the phylogenetic analysis presented in Figure 2 shows that both lineages 1 and 2 are embedded within clades containing sequences from CoV shed by *Chalinolobus gouldii*, the Australian sister species of *C. tuberculatus* (long-tailed bat). We further constructed a comprehensive phylogeny including 812 sequences that cover a substantial portion of the currently known bat-borne CoV diversity at a global scale. In this phylogeny (https://itol.embl.de/tree/92130128120274381648222266), lineage 1 is embedded within a clade containing viral sequences obtained from several bat species, including two distinct Australian *Chalinolobus* species, hence supporting a co-diversification of these CoV lineages and their *Chalinolobus* bat hosts from Australasia.

Of note, lesser short-tailed bats have been previously reported as hosting a CoV lineage (Hall et al., 2014), which was not detected in the present study. A detailed inspection of the RdRp sequence shows that the semi-nested PCR used herein is likely not adapted to the detection of this specific viral lineage, because of numerous mismatches in one of the used primers (CoV-Fwd2, not shown).

## Conclusions

From an epidemiological perspective, it is important to note that both reported lineages are embedded within alpha CoVb clade, while all three CoV that have emerged in human populations in the last decades are Beta-Cov, more specifically belonging to Sarbe- and Merbe-CoV subgenera. Therefore, the viral lineages presented herein cannot be classified as of highest risk of emergence in human populations.

From an evolutionary standpoint, the topologies of the two presented phylogenies are coherent with a *Chalinolobus* ancestral population(s) colonizing New Zealand from Australia while infected with CoV. Therefore, these viral sequences appear as good candidates to infer evolutionary rates of these viruses using molecular information from bat hosts, including mitochondrial sequences and microsatellites data (Dool et al., 2016). Note worthily, *Chalinolobus* is a relatively recent colonist of New Zealand (<1 myr), compared to lesser short-tailed bats (>20 my) (Lloyd, 2003; O’Donnell & Borkin, 2021), which is coherent with the limited diversity of associated CoV reported herein.

The high infection prevalence (reaching 61.7% in long-tailed bats), the dispersal capacities and the ability of New Zealand bats to enter into torpor make this biological model relevant to explore the mechanisms of maintenance of bat-borne CoVs in the wild. It has been proposed that torpor or hibernation are physiological adaptations that may facilitate viral maintenance in bats (Calisher et al., 2006). It is thus relevant to implement a longitudinal survey in order to address the temporal dynamics of these infections through one or two successive seasons. Such a non-invasive survey could allow identifying drivers of infection within colonies throughout the reproductive season (Dietrich et al., 2015; Joffrin et al., 2022) and provide novel information needed to identify the emergence drivers of these zoonotic viruses.

## Acknowledgements

We would like to thank Tania King from the University of Otago for invaluable technical advice as well as staff from the Department of Conservation of New Zealand for their contribution, especially Clément Lagrue for facilitating networking between all institutions involved in the study and Luke Easton for scat sample collection. This work was carried out in the frame of a sabbatical stay of Pablo Tortosa (Congés de Recherche et de Conversion Thématique) that was financed by the French Conseil National des Universités and by European Regional Development Fund/Regional Council of La Réunion (ERDF GeCaBex project).

## Notes

https://itol.embl.de/tree/92130128120274381648222266

### Competing Interest Statement

The authors have declared no competing interest.

## References

Anthony, S. J., Johnson, C. K., Greig, D. J., Kramer, S., Che, X., Wells, H., Hicks, A. L., Joly, D. O., Wolfe, N. D., Daszak, P., Karesh, W., Lipkin, W. I., Morse, S. S., PREDICT Consortium, Mazet, J. A. K., & Goldstein, T. (2017). Global patterns in coronavirus diversity. Virus Evolution, 3(1), vex012. https://doi.org/10.1093/ve/vex012

Calisher, C. H., Childs, J. E., Field, H. E., Holmes, K. V., & Schountz, T. (2006). Bats: Important reservoir hosts of emerging viruses. Clinical Microbiology Reviews, 19(3), 531–545. https://doi.org/10.1128/CMR.00017-06

Chan, J. F.-W., To, K. K.-W., Tse, H., Jin, D.-Y., & Yuen, K.-Y. (2013). Interspecies transmission and emergence of novel viruses: Lessons from bats and birds. Trends in Microbiology, 21(10), 544–555. https://doi.org/10.1016/j.tim.2013.05.005

Cui, J., Li, F., & Shi, Z.-L. (2019). Origin and evolution of pathogenic coronaviruses. Nature Reviews. Microbiology, 17(3), 181–192. https://doi.org/10.1038/s41579-018-0118-9

Darriba, D., Taboada, G. L., Doallo, R., & Posada, D. (2012). jModelTest 2: More models, new heuristics and parallel computing. Nature Methods, 9(8), 772–772. https://doi.org/10.1038/nmeth.2109

Dietrich, M., Wilkinson, D. A., Benlali, A., Lagadec, E., Ramasindrazana, B., Dellagi, K., & Tortosa, P. (2015). Leptospira and paramyxovirus infection dynamics in a bat maternity enlightens pathogen maintenance in wildlife. Environmental Microbiology, 17(11), 4280–4289. https://doi.org/10.1111/1462-2920.12766

Dool, S. E., O’Donnell, C. F. J., Monks, J. M., Puechmaille, S. J., & Kerth, G. (2016). Phylogeographic-based conservation implications for the New Zealand long-tailed bat, (Chalinolobus tuberculatus): Identification of a single ESU and a candidate population for genetic rescue. Conservation Genetics, 17(5), 1067–1079. https://doi.org/10.1007/s10592-016-0844-3

Drexler, J. F., Corman, V. M., Wegner, T., Tateno, A. F., Zerbinati, R. M., Gloza-Rausch, F., Seebens, A., Müller, M. A., & Drosten, C. (2011). Amplification of Emerging Viruses in a Bat Colony. Emerging Infectious Diseases, 17(3), 449–456. https://doi.org/10.3201/eid1703.100526

Hall, R. J., Wang, J., Todd, A. K., Bissielo, A. B., Yen, S., Strydom, H., Moore, N. E., Ren, X., Huang, Q. S., Carter, P. E., & Peacey, M. (2014). Evaluation of rapid and simple techniques for the enrichment of viruses prior to metagenomic virus discovery. Journal of Virological Methods, 195, 194–204. https://doi.org/10.1016/j.jviromet.2013.08.035

Joffrin, L., Goodman, S. M., Wilkinson, D. A., Ramasindrazana, B., Lagadec, E., Gomard, Y., Minter, G. L., Santos, A. D., Schoeman, M. C., Sookhareea, R., Tortosa, P., Julienne, S., Gudo, E. S., Mavingui, P., & Lebarbenchon, C. (2020). Bat coronavirus phylogeography in the Western Indian Ocean. Scientific Reports, 10(1), 1–11. https://doi.org/10.1038/s41598-020-63799-7

Joffrin, L., Hoarau, A. O. G., Lagadec, E., Torrontegi, O., Köster, M., Le Minter, G., Dietrich, M., Mavingui, P., & Lebarbenchon, C. (2022). Seasonality of coronavirus shedding in tropical bats. Royal Society Open Science, 9(2), 211600. https://doi.org/10.1098/rsos.211600

Lacroix, A., Duong, V., Hul, V., San, S., Davun, H., Omaliss, K., Chea, S., Hassanin, A., Theppangna, W., Silithammavong, S., Khammavong, K., Singhalath, S., Greatorex, Z., Fine, A. E., Goldstein, T., Olson, S., Joly, D. O., Keatts, L., Dussart, P., … Buchy, P. (2017). Genetic diversity of coronaviruses in bats in Lao PDR and Cambodia. Infection, Genetics and Evolution: Journal of Molecular Epidemiology and Evolutionary Genetics in Infectious Diseases, 48, 10–18. https://doi.org/10.1016/j.meegid.2016.11.029

Latinne, A., Hu, B., Olival, K. J., Zhu, G., Zhang, L., Li, H., Chmura, A. A., Field, H. E., Zambrana-Torrelio, C., Epstein, J. H., Li, B., Zhang, W., Wang, L.-F., Shi, Z.-L., & Daszak, P. (2020). Origin and cross-species transmission of bat coronaviruses in China. Nature Communications, 11(1), 4235. https://doi.org/10.1038/s41467-020-17687-3

Lemoine, F., Correia, D., Lefort, V., Doppelt-Azeroual, O., Mareuil, F., Cohen-Boulakia, S., & Gascuel, O. (2019). NGPhylogeny.fr: New generation phylogenetic services for non-specialists. Nucleic Acids Research, 47(W1), W260–W265. https://doi.org/10.1093/nar/gkz303

Lloyd, B. D. (2001). Advances in New Zealand mammalogy 1990–2000: Short-tailed bats. Journal of the Royal Society of New Zealand, 31(1), 59–81. https://doi.org/10.1080/03014223.2001.9517639

Lloyd, B. D. (2003). The demographic history of the New Zealand short-tailed bat Mystacina tuberculata inferred from modified control region sequences. Molecular Ecology, 12(7), 1895–1911. https://doi.org/10.1046/j.1365-294x.2003.01879.x

Maganga, G. D., Pinto, A., Mombo, I. M., Madjitobaye, M., Mbeang Beyeme, A. M., Boundenga, L., Ar Gouilh, M., N’Dilimabaka, N., Drexler, J. F., Drosten, C., & Leroy, E. M. (2020). Genetic diversity and ecology of coronaviruses hosted by cave-dwelling bats in Gabon. Scientific Reports, 10(1), 7314. https://doi.org/10.1038/s41598-020-64159-1

O’Donnell, C. F. J., Richter, S., Dool, S., Monks, J. M., & Kerth, G. (2016). Genetic diversity is maintained in the endangered New Zealand long-tailed bat (Chalinolobus tuberculatus) despite a closed social structure and regular population crashes. Conservation Genetics, 17(1), 91–102. https://doi.org/10.1007/s10592-015-0763-8

O’Donnell, C. F. J. (2000). Cryptic local populations in a temperate rainforest bat Chalinolobus tuberculatus in New Zealand. Animal Conservation, 3(4), 287–297. https://doi.org/10.1111/j.1469-1795.2000.tb00114.x

O’donnell, C. F. J. (2002). Variability in numbers of long-tailed bats (Chalinolobus tuberculatus) roosting in Grand Canyon Cave, New Zealand: Implications for monitoring population trends. New Zealand Journal of Zoology, 29(4), 273–284. https://doi.org/10.1080/03014223.2002.9518311

O’Donnell, C. F. J., Borkin, K., Christie, J., Lloyd, B., Parsons, S., & Hitchmough, R. (2018). Conservation status of New Zealand bats, 2017 (New Zealand Threat Classification Series, 21). New Zealand Department of Conservation. https://www.doc.govt.nz/globalassets/documents/science-and-technical/nztcs21.pdf

O’Donnell, C. F. J., & Borkin, K. M. (2021). Chalinolobus tuberculatus. In The Handbook of New Zealand Mammals (CM King&DM Forsyth, pp. 95–130). CSIRO Publishing.

O’Donnell, C. F. J., Christie, J. E., Hitchmough, R. A., Lloyd, B., & Parsons, S. (2010). The conservation status of New Zealand bats, 2009. New Zealand Journal of Zoology, 37(4), 297–311. https://doi.org/10.1080/03014223.2010.513395

O’Donnell, C. F. J., & Sedgeley, J. A. (2006). Causes and consequences of tree-cavity roosting in a temperate bat, Chalinolobus tuberculatus, from New Zealand. In Functional and Evolutionary Biology of Bats (Zubaid, A., McCracken, G. F. & Kunz, T. H., pp. 308–328). Oxford University Press.

Peel, A. J., Field, H. E., Aravena, M. R., Edson, D., McCallum, H., Plowright, R. K., & Prada, D. (2020). Coronaviruses and Australian bats: A review in the midst of a pandemic. Australian Journal of Zoology. https://doi.org/10.1071/ZO20046

Poon, L. L. M., Chu, D. K. W., Chan, K. H., Wong, O. K., Ellis, T. M., Leung, Y. H. C., Lau, S. K. P., Woo, P. C. Y., Suen, K. Y., Yuen, K. Y., Guan, Y., & Peiris, J. S. M. (2005). Identification of a novel coronavirus in bats. Journal of Virology, 79(4), 2001–2009. https://doi.org/10.1128/JVI.79.4.2001-2009.2005

Prada, D., Boyd, V., Baker, M. L., O’Dea, M., & Jackson, B. (2019). Viral Diversity of Microbats within the South West Botanical Province of Western Australia. Viruses, 11(12). https://doi.org/10.3390/v11121157

Rambaut, A. (2014). FigTree 1.4.2 software. Institute of Evolutionary Biology, University of Edinburgh. http://tree.bio.ed.ac.uk/software/figtree/

Ronquist, F., Teslenko, M., van der Mark, P., Ayres, D. L., Darling, A., Hohna, S., Larget, B., Liu, L., Suchard, M. A., & Huelsenbeck, J. P. (2012). MrBayes 3.2: Efficient Bayesian phylogenetic inference and model choice across a large model space. Systematic Biology, 61(3), 539–542. https://doi.org/10.1093/sysbio/sys029

Smith, C. S., de Jong, C. E., Meers, J., Henning, J., Wang, L.-F., & Field, H. E. (2016). Coronavirus Infection and Diversity in Bats in the Australasian Region. EcoHealth, 13(1), 72–82. https://doi.org/10.1007/s10393-016-1116-x

Tortosa, P., Pascalis, H., Guernier, V., Cardinale, E., Le Corre, M., Goodman, S. M., & Dellagi, K. (2012). Deciphering arboviral emergence within insular ecosystems. Infection, Genetics and Evolution, 12(6), 1333–1339. https://doi.org/10.1016/j.meegid.2012.03.024

Wacharapluesadee, S., Duengkae, P., Chaiyes, A., Kaewpom, T., Rodpan, A., Yingsakmongkon, S., Petcharat, S., Phengsakul, P., Maneeorn, P., & Hemachudha, T. (2018). Longitudinal study of age-specific pattern of coronavirus infection in Lyle’s flying fox (Pteropus lylei) in Thailand. Virology Journal, 15(1), 38. https://doi.org/10.1186/s12985-018-0950-6

Wang, J., Moore, N. E., Murray, Z. L., McInnes, K., White, D. J., Tompkins, D. M., & Hall, R. J. (2015). Discovery of novel virus sequences in an isolated and threatened bat species, the New Zealand lesser short-tailed bat (Mystacina tuberculata). The Journal of General Virology, 96(8), 2442–2452. https://doi.org/10.1099/vir.0.000158

